# Recombinant *Helicobacter pylori* CagA protein induces endoplasmic reticulum stress and autophagy in human cells

**DOI:** 10.1101/2020.06.26.174615

**Authors:** Babak Nami, Ali Azzawri, Vasfiye B Ucar, Hasan Acar

## Abstract

*Helicobacter pylori* (*Hp*) CagA protein has a key role in the development of gastric cancer by the intruding in many intracellular processes of host human cell. Endoplasmic reticulum (ER) stress is an essential process for cellular homeostasis that modulates survival and death and is linked to several complex diseases including cancer. CagA protein is found in the serum of *Hp*-positive individuals and also in the supernatant of *Hp* culture. Limited studies report that recombinant CagA can alter gene expression and signaling pathways and induce the death of human cells. In this study, we investigated the effect of exogenous recombinant CagA protein treatment on ER stress and autophagy of human cell. AGS, MKN45, and HEK293 cells were treated with 1 µg/ml of recombinant CagA protein and then ER stress was studied by quantitative-PCR of spliced XBP-1 mRNA, immunofluorescence staining of ATF6 protein nuclear localization and real-time quantitative-PCR and/or western blot expression of GRP78, GRP94, ATF4 and CHOP genes. Autophagy was studied by western blot assessment of the conversion of LC3-I to LC3-II and LC3 aggregation. Cell proliferation and death were investigated by MTT assay and trypan blue staining respectively. As result, treatment with recombinant CagA enhanced XBP-1mRNA splicing, nuclear localization of ATF6, and the expression of ER stress signaling target genes in the cells. Recombinant CagA also induced LC3 protein conversion and aggregation in the cells. Reduced cell proliferation and increased cell death were determined in the cells treated with recombinant CagA. These results show that exogenous recombinant CagA protein causes cell death by inducing ER stress and autophagy in human cells. We conclude that CagA protein exogenously localizes in/on human cells and induces ER stress via disturbing protein machinery leading the human cell to death, however, the mechanism of CagA-host cell interaction is to be investigated.

## Introduction

*Helicobacter pylori* (*Hp*) is a gram-negative and microaerophilic bacterium that is found in the stomach of more than half of people [1]. *Hp* is associated with many stomach diseases like gastritis, peptic, duodenal ulcers, and gastric adenocarcinoma [2]. Approximately, 90% of people with duodenal ulcers, and 80% of patients with gastric cancer were the host for *Hp* [3]. Virulence factors of *Hp* comprise a wide range of structural elements like outer proteins and secretary enzymes particularly exotoxins including cytotoxin associated gene A (CagA) [4]. The genome of the most virulent strains associated with a higher risk of gastric adenocarcinoma possesses a 40 kbp DNA stretch, harbor in their genome cluster of 31 genes called Cag pathogenicity island (cagPAI) encoding a type IV secretion system (T4SS) and CagA gene [5]. T4SS is a needle-like structure protruding from the *Hp* surface that interacts with the host cell and injects the CagA protein into the cytoplasm of the host cell [4,5]. CagA gene encodes about a 1190 amino acid protein with a molecular weight of 120-145 kDa [6]. After the injection of CagA into the host cell, it can be phosphorylated from the pentamerous amino acid sequence called EPIYA (glutamic acid, proline, isoleucine, tyrosine, and Alanine) motifs in C-terminus residue, by the host cell Src and Abl kinases and turn to an active form [6,7].

The endoplasmic reticulum (ER) is a labyrinth-like primary organelle that is involved in the synthesis, folding, arraignment, and trafficking of proteins and is also a membrane factory and pool of calcium ion for eukaryotic cells [8]. Unfolded/misfolded proteins in eukaryotic cells potentially are toxic and must be folded or deleted, otherwise, they can accumulate in ER lumen [9]. Accumulation of faulty proteins in ER leads to unfolded protein response (UPR) and ER stress which follow two primary aims: the first is to restore normal cellular function by halting protein translation and synthesis [10–12], and the second aim is to activate ER stress signaling that leads to increase the production of molecular chaperones involved in protein folding and degradation [13]. If these aims are not achieved within a certain time, the ER stress leads to activation programmed cell death via apoptosis and autophagy [14]. Therefore, ER stress is in charge of guarding the intracellular homeostasis by haltering cell survival and death mechanisms. ER stress plays a pivotal role in the development of inflammatory disorders, obesity, diabetes, and cancer and is responsible for neural cell death in neurodegenerative diseases [15].

ER stress signaling pathway is framed with three distinct branches which can be promoted by three transmembrane proteins anchored in ER membrane with their N-terminus in the lumen of the ER and their C-terminus in the cytosol, including PRKR-like ER kinase (PERK) which is also known as EIF2AK3, inositol-requiring kinase 1α (IRE1α) and activating transcription factor 6 (ATF6) [11,14]. When unfolded/misfolded protein accumulation occurs, GRP78, a heat shock protein family member chaperon, releases from N-terminus of these transmembrane proteins, allowing their oligomerization and thereby initiating the downstream cascade [14,15]. IRE1α is a type I transmembrane protein that has both a serine/threonine protein kinase and an endoribonuclease domain [16]. Activated IRE1α binds to TRAF2 that upregulates JNK, BCL-2, P38 MAPK, and eventually caspases, the essential compounds for initiation of apoptosis [17]. IRE1α also brings out a 26 bp intron from X-box-binding protein-1 (XBP-1) pre-mRNA and splices it to mature mRNA using its endoribonuclease domain [18]. Splicing of XBP-1 RNA allows the synthesis of XBP-1 protein, a basic leucine zipper (bZIP) family transcription factor that can increase the transcription of chaperons and endoplasmic reticulum-associated degradation (ERAD) genes, also CHOP a pro-apoptotic protein [18,19]. PERK is another sensor that is charged to phosphorylate eIF2α. Phosphorylated elF2α has two main functions: the suspension of global protein synthesis and increasing ATF4 protein production [20]. ATF4 is a member of bZIP family of transcription factors that upregulates the gene expression of ER chaperones as well as the genes involved in apoptosis and autophagy [21]. ATF6 is another transmembrane sensor that is functionally similar to ATF4. When ER stress is activated, ATF6 releases from ER and immigrates to Golgi apparatus wherein can be cleaved into P90 and P50 fragments. The N-terminus P50 fragment is a transcription factor, which translocates to the nucleus and regulates the gene expression [22].

The role of ER stress in tumor promotion has been well-demonstrated. Overexpression of GRP78 in many types of tumors has been reported. GRP78 has been suggested as an oncogene and a potential target for cancer therapy [23,24]. Several studies have implicated that the IRE1α/XBP-1 arm of ER stress signaling regulates expression of the genes involved in inflammation, angiogenesis, and tumorigenesis [25]. More studies have also suggested that XBP-1 may be a biomarker and target for tumor diagnosis and therapy [26]. Besides, a significant role of PERK/eIF2α pathway on carcinogenesis has been reported [27]. It has been indicated that ATF4 has associated with tumor neoplasia as well as cancer chemoresistance [28]. We have shown that drug-induced ER stress in the breast cancer cell culture high expression of ATF4 and CHOP reduces subpopulation of cancer stem cells inside the bulk cells, which results in reduced in vitro invasion [29]. Also, we reported a positive correlation between the expression of molecular chaperons GRP78 and GRP94 and the emergence of breast cancer stem cell phenotype [24]. These show that interstice upregulation of ER stress elements increases cancer cell survival, while induction of ER stress suppresses cancer cell invasion by reducing the cancer stem cell subpopulation.

It has been demonstrated that CagA protein is tightly associated with gastric cancer development [7], however, little is known about the effect of CagA in ER stress of the host cell. Many studies on the effect of CagA on mammalian cells were carried out by transgene expression of CagA, or co-culturing of cells with CagA-positive *Hp* [7]. However, some other studies showed that recombinant CagA protein altered gene expression, signal pathways, and induced apoptosis in human cells. In addition, CagA protein was found in the serum of *Hp-* positive individuals [30,31] as well as in the supernatant of in vitro *Hp* culture [32,33]. These pieces of evidence suggest that not only intracellular CagA, but also extracellular exogenous CagA in the microenvironment of human cells affects the physiobiological activities of the cells [34–37]. In this study, we aimed to investigate the effect of exogenous recombinant CagA on the ER stress signaling and autophagy of human cells.

## Results

### Recombinant CagA induces XBP-1 mRNA splicing

To investigate whether recombinant CagA induces ER stress, we treated AGS, MKN45, and HEK293 cells with 1 µg/ml CagA for 2 and 4 hours and then analyzed XBP-1 mRNA splicing. A final concentration of 2 µg/ml of tunicamycin and untreated were used as respectively, positive and mock controls. Splicing of XBP-1 mRNA is a specific marker for ER stress that is indicated by the deletion of a 26 bp intron sequence by active IRE1α [18]. By using the same primer pair for XBP-1 cDNA PCR, we have previously shown that 289 bp amplicon represents an unspliced (pre-mRNA) and 263 bp amplicon represents a spliced (mature mRNA) XBP-1 mRNA [29]. Results showed an increased level of spliced XBP-1 in the cells treated with CagA in comparison with mock control. The cells treated with the combination of CagA and tunicamycin showed a significantly higher level of spliced XBP-1 than it in the control cells (Fig. 1B). These results indicate that recombinant CagA enhances XBP-1 mRNA splicing, indicating that recombinant CagA induces ER stress in the cells.

**Figure 1.**
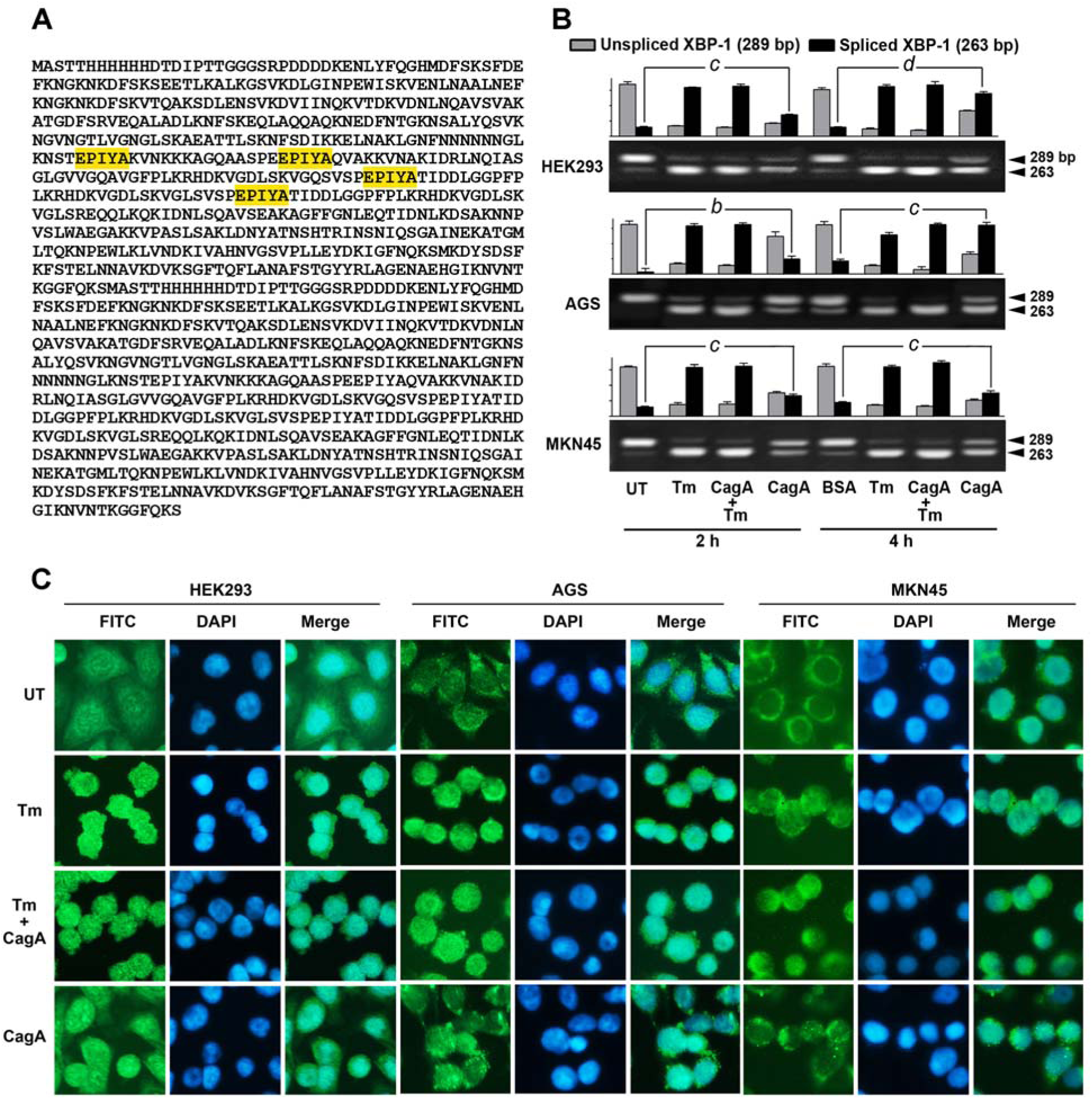
Recombinant CagA induces XBP-1 mRNA splicing. **(A)** The amino acid sequence of C-terminus CagA. EPIYA motives are highlighted by yellow. **(B)** Quantitative PCR result showing XBP-1 mRNA level in HEK293, AGS and MKN45 cells treated with 1 µg/ml of exogenous CagA or for 2 and 4 hours. The panel shows SDS-PAGE band and band intensity of spliced (263 bp) and unspliced (289 bp) XBP-1 mRNA amplicons amplified from cDNA using the same primer pairs and. Two µg/ml of tunicamycin (Tm) was used as positive control. *b*; *P* ≤ 0.01, *c*; *P* ≤ 0.001, *d*; *P* ≤ 0.0001. (**C**) Immunofluorescence staining of cleaved ATF6 protein (FITC; green) in HEK293, AGS, and MKN45 cells treated with 1 µg/ml of exogenous CagA for 4 hours. Nuclei were mounted by DAPI (Blue). Two µg/ml of tunicamycin (Tm) was used as positive control. The size of each side of the small image square is equal to approximately 100 µm.

### Recombinant CagA induces ATF6 nuclear translocation

Translocation of ATF6 protein from the ER to the nucleus is a hallmark for activated ER stress [22]. To investigate whether exogenous recombinant CagA induces ATF6 nuclear translocation, we treated the cells with 1 µg/ml CagA for 4 hours and the investigated the localization of ATF6 by immunofluorescence staining using an antibody against C-terminal end of ATF6 protein. As shown in Fig. 1C, a significant ATF6 localization was detected in the nucleus of the cells treated with CagA but not in the mock control cells (Fig. 1C). Also, strong nuclear ATF6 was detected in the cells treated with tunicamycin as well as in those treated with a combination of tunicamycin and CagA. These results indicate that recombinant CagA induces ATF6 nuclear translocation in the cells.

### Recombinant CagA upregulates ER stress target genes

For further confirmation, we investigated the transcriptional and translational expression levels of ER stress downstream target genes in response to exogenous recombinant CagA treatment. For this, the cells were treated with 1 µg/ml CagA for 2 and 4 hours, and then mRNA expression of GRP78, GRP94, ATF4, and CHOP was investigated by real-time qPCR. Also, the protein expression of ATF4 and CHOP was studied by western blotting. As result, CagA significantly increased the transcription of GRP78, GRP94, ATF4, and CHOP in the three cells in 2- and 4-hours treatments (Fig. 2A). Further, increased levels of GRP78 and ATF4 proteins were detected in the cells in response to recombinant CagA (Fig. 2B). Taken together, these results show that recombinant CagA upregulates the expression of ER stress target genes.

**Figure 2.**
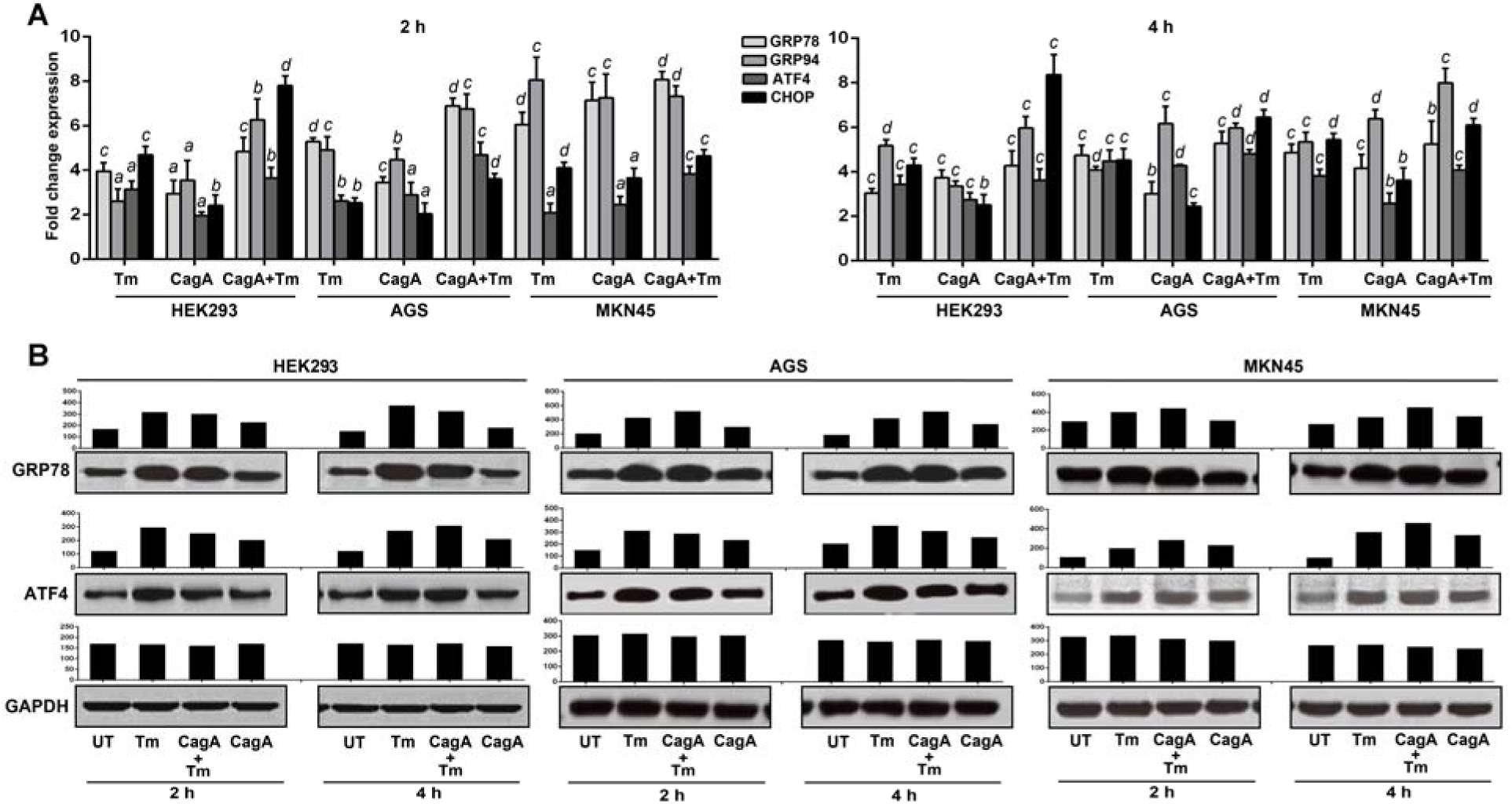
Recombinant CagA upregulates the expression of ER stress targets. **(A)** Quantitative Real-time PCR mRNA expression level of GRP78, GRP94, ATF4 and CHOP in HEK293, AGS and MKN45 cells treated 1 µg/ml of exogenous or CagA for 2 and 4 hours. **(B)** Western blot protein expression of GRP78 and ATF4 in the cells treated with CagA. Two µg/ml of tunicamycin (Tm) was used as positive control. GAPDH was used as housekeeping gene expression and loading controls. *a*; *P* ≤ 0.05, *b*; *P* ≤ 0.01, *c*; *P* ≤ 0.001, *d*; *P* ≤ 0.0001.

### Recombinant CagA induces autophagy

ER stress is a mechanism of autophagy initiation. To examine whether exogenous recombinant CagA treatment induces autophagy in the host cells, the cells were treated with 1 µg/ml CagA for 24, 48, and 72 hours and then LC3-I to LC3-II conversion was studied by western blotting. In addition, LC3 protein aggregation was investigated by immunofluorescence staining. Treatment with 20 µM of C2-ceramide was used as autophagy inducer positive control. Untreated cells were used as mock control. The result showed an increased level of LC3-I to LC3-II conversion (Fig 3A), and LC3 aggregation (Fig 3B and C) in the cells treated with recombinant CagA. These results indicate that recombinant CagA increases LC3 protein conversion and autophagosome formation showing that exogenous recombinant CagA induces autophagy of the host cells.

**Figure 3.**
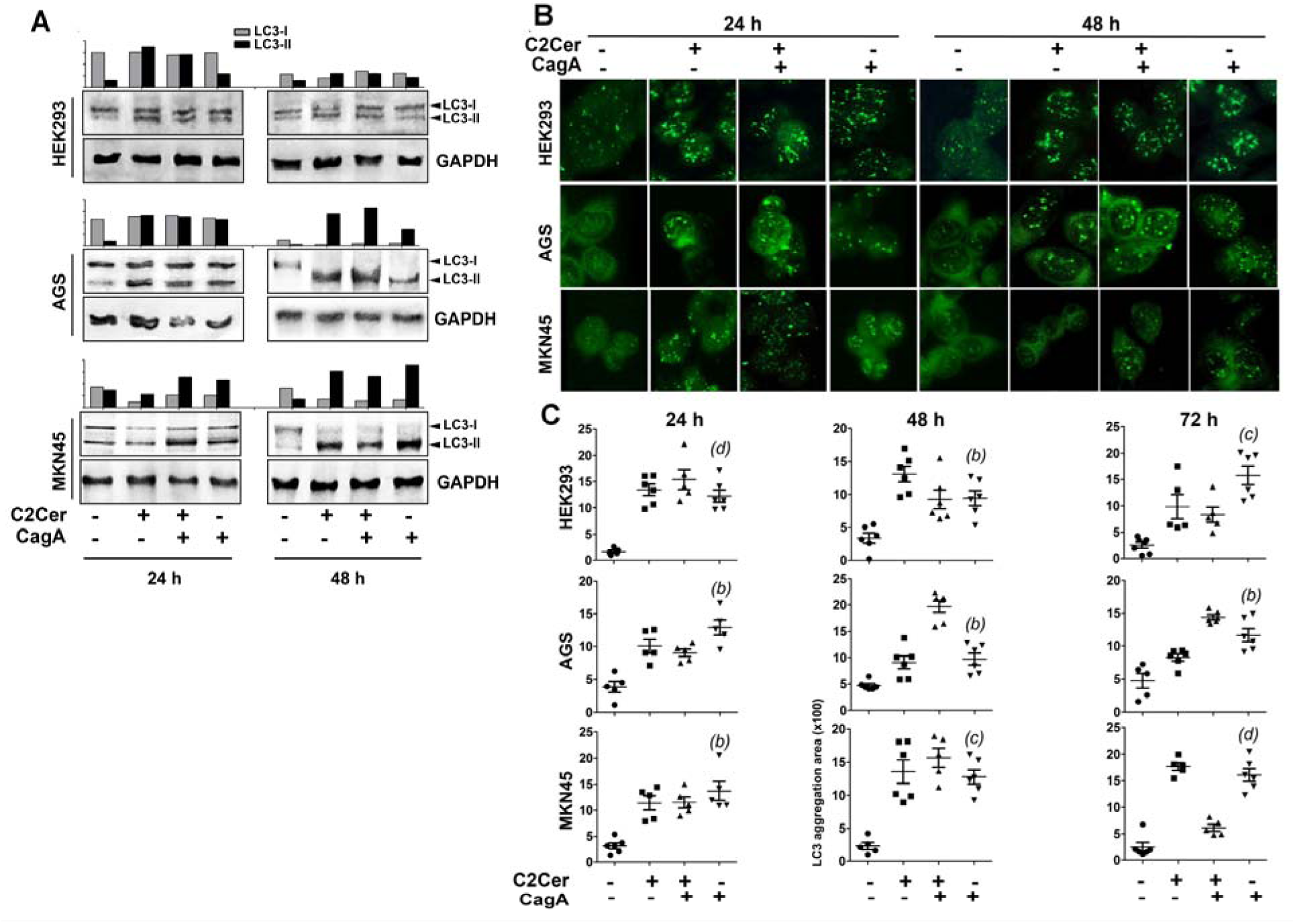
Recombinant CagA induces LC3 conversion and aggregation. HEK293, AGS, and MKN45 cells were treated with 1 µg/ml of exogenous CagA or 20 µM of C2-ceramide (C2Cer) as positive control for autophagy for 24 and 48 and 72 hours. **(A)** Western blot results of LC3-I and LC3-II protein isoforms. GAPDH was used as loading control. **(B)** Immunofluorescence staining of LC3 proteins (FITC; green) showing autophagosomes. (**C)** Quantitative levels of LC3 aggregation spots (autophagosomes) in 5 cells from each treated cell group which were analyzed by LC3 immunofluorescence staining (see panel **B**). The size of each side of the small image square is equal to approximately 100 µm. *a*; *P* ≤ 0.05, *b*; *P* ≤ 0.01, *c*; *P* ≤ 0.001, *d*; *P* ≤ 0.0001.

### Recombinant CagA inhibits cell proliferation via inducing cell death

Our result showed that CagA induces ER stress-mediated autophagy in the cells. Autophagy is a *bona fide* mechanism of cell death. To examine whether recombinant CagA suppresses cell proliferation by induction cell death, we treated the cells with 1 µg/ml recombinant CagA for 24, 48, and 72 hours, then investigated cell proliferation and death. Results showed a significant decline in the proliferation of the cell lines in response to the CagA treatment compared to untreated cells (Fig. 4A). Trypan blue staining of HEK293 cells treated with 1 µg/ml recombinant CagA for 48 hours showed a dramatically increased cell death compared to the untreated cells (Fig. 4B). Taken together, these results show that treatment of human cells with exogenous recombinant CagA induces human cell death.

**Figure 4.**
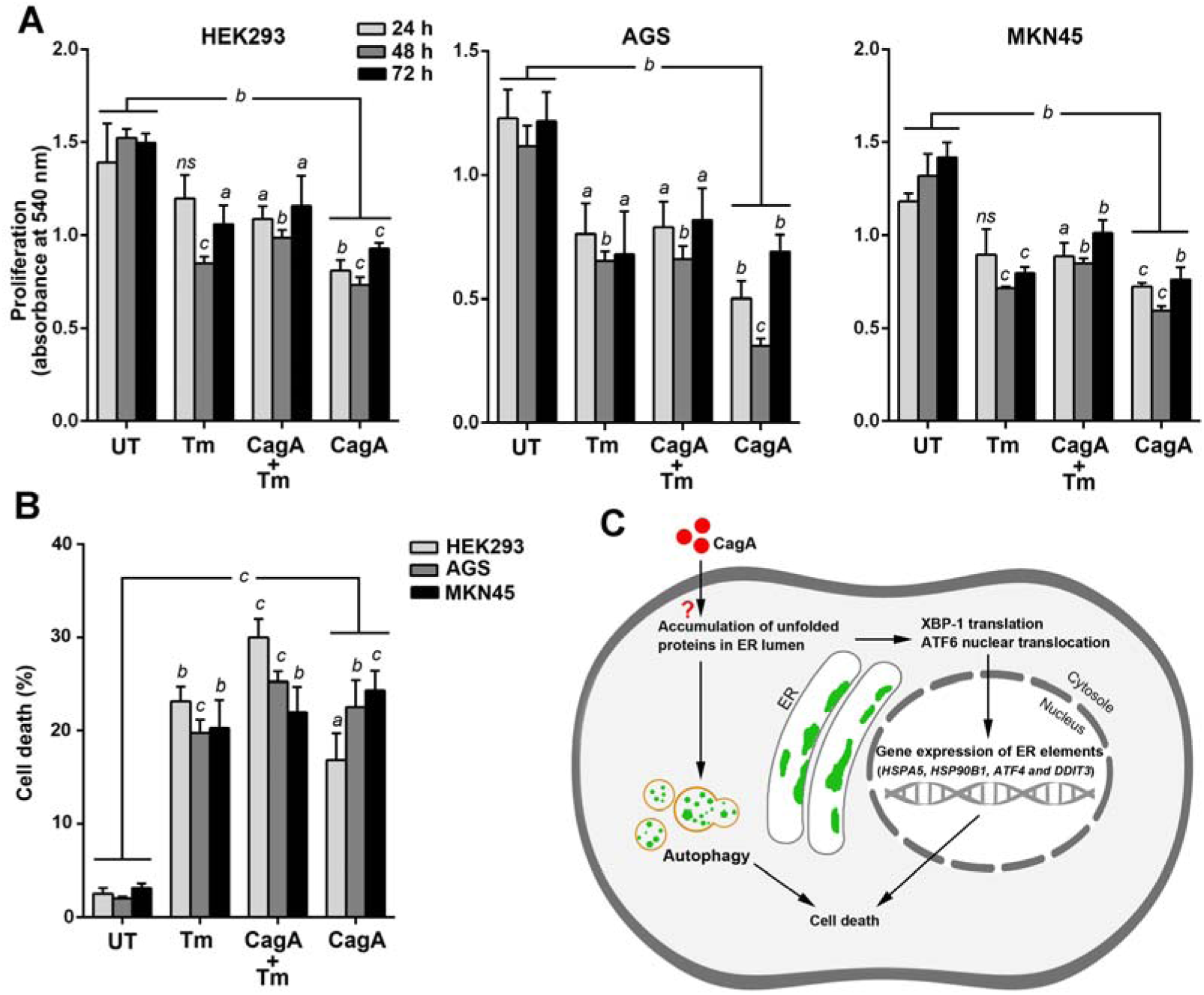
Recombinant CagA induces cell death in human cells. HEK293, AGS, and MKN45 cells were treated with 1 µg/ml of exogenous CagA for 24, 48, and 72 hours. **(A)** Cell proliferation levels of the cells determined by MTT assay. **(B)** The cell death rate of HEK293 cells determined by trypan blue staining. Two µg/ml of tunicamycin (Tm) was used as positive control. GAPDH was used as housekeeping control gene. *a*; *P* ≤ 0.05, *b*; *p* ≤ 0.01, *c*; *P* ≤ 0.001, *d*; *P* ≤ 0.0001, ns; Not significant. **(C)** Model summarizing the effect of exogenous CagA protein on ER stress and autophagy of human cell. The question mark refers to an unknown mode of interaction of extracellular CagA with human cells. CagA causes accumulation of unfolded proteins in the ER lumen and induces unfolded protein response and autophagy in the cell. During ER stress mRNA of XBP-1 is spliced by the endoribonuclease domain of the ER transmembrane protein IRE1 and XBP-1 protein is synthesized. ATF6 releases from the ER membrane and localizes in the nucleus. Both XBP-1 and ATF6 are b-zipper transcription factors that promote the expression of ER genes. Induction of ER stress and autophagy lead the cell to death.

## Discussion

In this study, we investigated the effect of exogenous recombinant C-terminal part of *Hp* CagA protein on the ER stress and autophagy. We here used the C-terminal part of CagA for two main reasons; (1) C-terminus part of CagA consists EPIYA motifs, the sequences that are essential for phosphorylation of CagA by host cell kinases and is known essential sequences for CagA bioactivity [5], and (2) full-length CagA is a large protein and is hard to produce. Our results showed that recombinant CagA significantly upregulated XBP-1 mRNA splicing and nuclear translocation of ATF6. It also increased the expression of ER stress target genes including ER chaperons GRP78 and GRP94 as well as the proapoptotic transcription factors ATF4 and CHOP. Our results implicate that treatment with exogenous recombinant CagA triggers ER stress in both cancerous AGS and MKN45 cells and also noncancerous HEK293 cells. ER stress is the preserver of the balance between cell survival and death when protein processing is impaired. According to the previous studies, activation of ER stress may lead cells to survival by upregulation of ER chaperones. Overactivation of the ER chaperones GRP78 and GRP94 in tumors [38] and tumor stem cells [24] has been demonstrated. Several reports are implicating the significant role of XBP-1 [26], ATF4 [27], and ATF6 [28] transcription factors in survival, invasion, and chemoresistance of many tumor types. These mediators consist of NFY DNA binding domains with the ability of binding to the ERSE DNA sequences of ERAD genes, a group of genes that are necessary for regulation of protein synthesis and maintaining cell homeostasis.

Our results also show that recombinant CagA treatment induces the formation of autophagosomes in the host cells. Autophagy has a key role in the determination of cancer fate [39]. We previously showed a dramatic decline in the cancer stem cell population in culture under the autophagic condition [40]. Since autophagy is a consequence of active ER stress, the promotion of autophagy by recombinant CagA might attribute to its activity to induce ER stress. Moreover, the results show an increased rate of cell death and inhibited proliferation due to CagA treatment. Programmed cell death can be set up with the formation of ER stress-mediated autophagy [39]. It has been demonstrated that when ER stress prolongs to a certain period, apoptosis and autophagy being to be activated [41]. Moreover, two downstream target modulators of ER stress, ATF4, and CHOP, have also a proapoptotic activity that promotes programmed cell death via apoptosis and autophagy [29,42]. Therefore, we conclude that ER stress-mediated autophagy might be the mechanism of host cell death by CagA.

Secreted CagA protein has been found in the serologic fluid of *Hp*-positive individuals [30,31] as well as in the supernatant of *Hp* culture medium [32,33]. Therefore, CagA protein can be found in the extracellular matrix and directly interact with host human cells. Limited studies suggest that recombinant CagA can alter cell signaling activity, gene expression, and can induce cell death [34–37]. Targosz et al. [34] treated the cells with a high level of exogenous CagA (10 µg/ml) while we used only 1 µg/ml of the protein for treatment in the present study. They showed that exogenous CagA proteins promoted apoptosis by up-regulation of *Cox-2* gene in MKN45 cells. Yang et al. [37] showed that treatment of mouse podocyte cells with recombinant CagA protein decreased the expression and membrane distribution of tight junction protein ZO-1, disrupted the filtration barrier function and increased the activation of p38 MAPK signaling pathway in the cells. Taken together, the results of very significant activity in interaction with human cells.

Accumulation of unfolded/misfolded proteins in the ER lumen promotes ER stress signaling and autophagy.[10] Effect of CagA treatment in inducing ER stress can be explained either CagA itself is accumulate in the ER lumen or it interacts with a set of endogenous host cell factors that trigger ER stress. In both mechanisms, extracellular CagA proteins need to enter the host cell. Yang et al., [37] suggest the exogenous recombinant CagA can enter the cells via endocytosis. However, there is no report showing the entry mechanism of extracellular CagA into the host cell in the absence of *Hp* T4SS. We here suggest four possible mechanisms for the extracellular CagA influx. (1) Via penetration; which refers to direct diffusion of a particle across the lipid bi-layer plasma membrane. This ability depends on certain situations including particle composition, size, shape, charge, hydrophobicity, and molecular dynamic of the plasma membrane. Penetration is the main method by which cells uptakes cell-penetrating peptides (CPPs) which are short peptides with the ability to inject themselves into cells directly through the plasma membrane. These peptides are usually highly cationic and rich in arginine and lysine amino acids [43]. However, C-terminus CagA used in this study is a slightly cationic protein in which the percentage of arginine and lysine amino acids in its structure are 1.6% and 11.1%. respectively. (2) By membrane transports; in this mechanism CagA protein may use membrane transporter proteins that permit flux of CagA across the plasma membrane. Currently, membrane transporters are used for the delivery of therapeutic agents into cells [44]. (3) Endocytosis; which is an active transportation mechanism by which a cell can devour large molecules including proteins. It occurs when a molecule attaches to cell membrane or a receptor on it, the membrane starts wrapping up the molecule resulting in the formation of an intracellular vesicle by invagination of the plasma membrane and membrane fusion until the protein is fully inside [45]. This mechanism is the most possible mechanism of extracellular CagA protein entry. In another mechanism is the binding of extracellular CagA to host cell receptor(s). In this mechanism presence of CagA inside the cell is not necessary. CagA may show ligand-like activity that binding to target cell receptor(s) activates downstream signaling leading to induction of ER stress. Interestingly, it has been demonstrated that ER stress can be activated via a series of cell surface receptors [46,47]. For example stimulation of β-adrenergic receptors which belong to the family of G protein-coupled receptors, alters ER calcium ion homeostasis by elevating intracellular calcium concentration that causes ER stress-induced apoptotic cell death via activating caspase-12 [48,49]. Other examples of ER stress inducer surface receptors are androgen (AR) and glucocorticoid (GR) receptors which are nuclear receptors. Segawa et al [50] showed that androgenic hormone-mediated activation of androgen receptor induces ER stress signaling in prostate cancer cells. In this case, androgen receptor binding to gene regulatory target sites activates the IRE1a and simultaneously inhibits PERK signaling leading to ER stress [51]. GR can disrupt protein machinery by regulating protein chaperons particularly heat shock proteins 70 and 90 and altering protein folding [52]. Also, several studies demonstrated that GR protects cells from ER stress-induced apoptosis via ameliorating ER stress by promoting the correct folding of proteins and enhancing the removal of misfolded proteins from the ER [53,54].

In conclusion, the mechanism by which recombinant CagA affects protein machinery and induces ER stress and autophagy need to be addressed. we strongly suggest investigating the exact mode of interaction of extracellular CagA protein with human cells to clarify molecular mechanism CagA in induction of host cell ER stress and autophagy. ER stress is an important cellular phenomenon that protects cellular homeostasis by equilibration of survival and death. ER stress is rendered to be a suitable target for cancer therapy in recent years. We previously reported that drug-induced ER stress featured with upregulation of ATF4 and CHOP induces MCF7 cell death and blocks in vitro invasion by decreasing cancer stem cell subpopulation [29]. Bortezomib (Valcade®) is an ER stress inducer drug approved by the FDA for the treatment of multiple myeloma. This compound can lead tumor cells to ER stress-associated death by inhibiting proteasomes. Eeyarestatin I is another ER targeting drug that can block the ERAD family of genes thereby induces death in tumor cells [55]. Therefore, we suggest testing the antitumor efficiency of bortezomib and eeyarestatin I in *Hp*-positive patients with gastric adenocarcinoma.

## Materials and methods

### CagA and chemicals

Recombinant C-terminal part of CagA protein (residues 61-581) containing a 6x histidine tag was purchased from Bioclone Inc. (San Diego, USA; #PQ0129). The amino acid sequence of used CagA protein is shown in Fig. 1A. CagA protein was dissolved in PBS. Tunicamycin and C2-ceramide (N-Acetyl-D-sphingosine) were obtained from Sigma-Aldrich (St. Luis, USA) and were dissolved in dimethyl sulfoxide (DMSO) and stored at +4°C.

### Cell culture and treatments

Human gastric cancer AGS (#C10071) and MKN45 (#C10137) and human embryonic kidney HEK293 (#C10139) cell lines were purchased from the National Cell Bank of Iran (NCBI; Tehran, Iran). To perform the experiments, the cells were seeded as 10^4^, 5×10^4^, 10^5^ and 5×10^5^ cells in 96, 24, 12 and 6-well plates respectively and cultured in Dulbecco’s Modified Eagle’s Medium (DMEM) containing 10% fetal bovine serum, 1% L-glutamine, 1% penicillin (100 unit/ml) and 1% streptomycin (10 mg/ml; all were from Gibco, Rockford, USA) in a 5% CO2 at 37°C condition. After the cells reached 80% confluency, the cells were harvested and precultured for treatments. CagA was dissolved in 6 M urea as 1µg/µl concentration and added to complete culture medium to a final concentration of 1µg/ml for treatment. Treatments were done for 1, 2, 4, 24, 48, and 72 h. A concentration of 2 µg/ml of tunicamycin and 20 µM C2-ceramide were used as respectively, positive control for ER stress autophagy inducers respectively.

### XBP-1 mRNA splicing assay

XBP-1 mRNA splicing assay was done by PCR as described previously [29] Used primer sequences were obtained from a previous study [29] and are shown in Table 1.

**Table 1.**
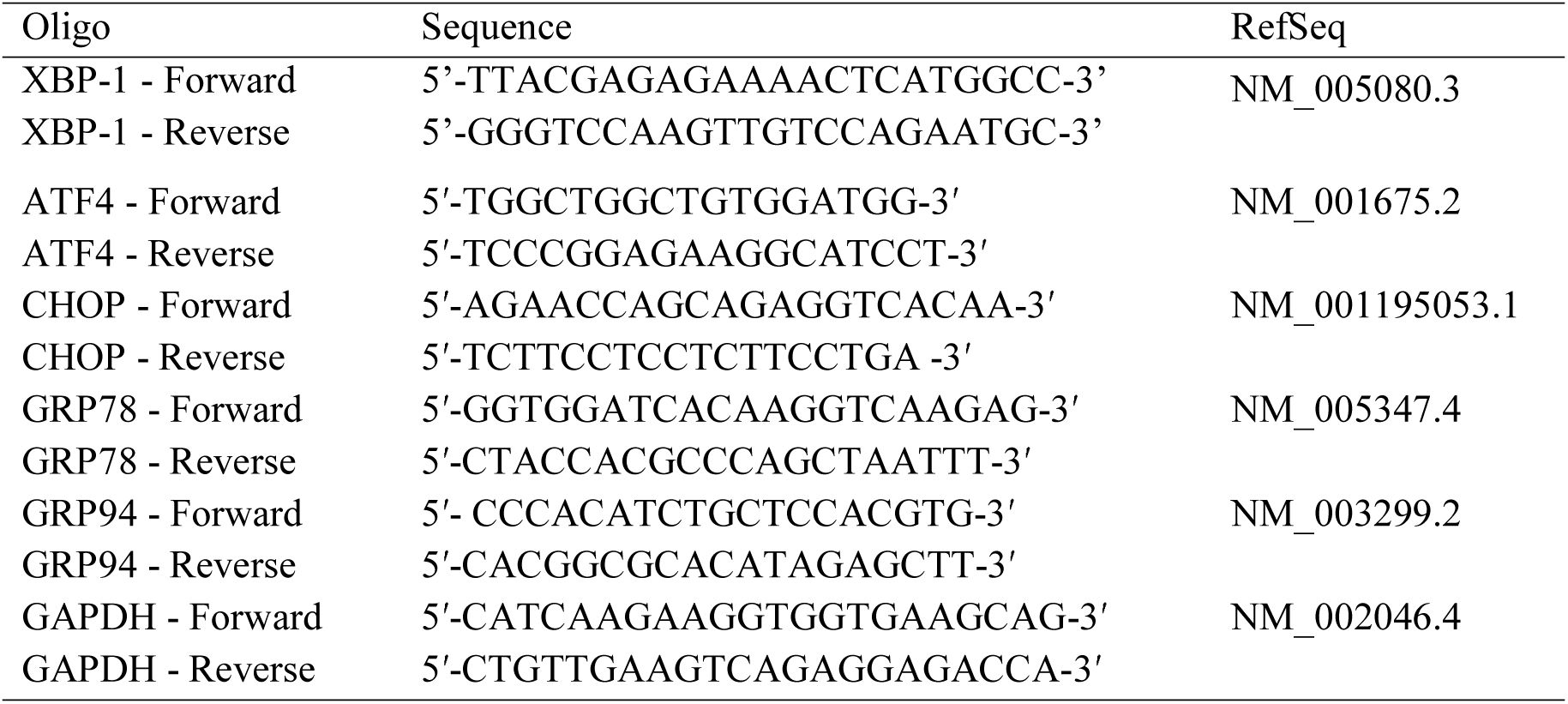
The sequences of the PCR primer oligos used in qPCR and their RefSeq numbers.

### DNA and protein polyacrylamide gel electrophoresis

Amount 2 µg of cDNA was mixed with 4 µl of 6x sample loading dye and ran in 10% polyacrylamide gel in 40 mA electric current for 1 hour. The gel then was stained in 1 µg/ml ethidium bromide solution. Total protein was extracted as described previously [29]. Amount 40 µg of total protein was run in 10% polyacrylamide gel at the same electrophoresis condition and then the gel was stained in coomassie blue solution (Bio-Rad Laboratories, Berkeley, USA; #1610786).

### Immunofluorescence staining assay

Immunofluorescence staining assay was carried out as described previously [29,40]

### RNA isolation and real-time qPCR

Total RNA from was extracted using TRIzol® reagent (Invitrogen, Waltham, USA) according to the protocol described previously [56]. cDNA synthesis was done by Transcriptor High-Fidelity cDNA Synthesis kit (Roche, Basel, Switzerland), using the oligo(dT)18 primer pairs following the manufacturer’s instructions. Oligonucleotide primers were synthesized by Biomers Inc. (Ulm, Germany). All primers used are listed in Table 1. All PCR reactions were carried out using SYBR Green/ROX qPCR Master Mix (Fermentas, Thermo Scientific, Rockford, USA) by Rotor-Gene-Q (Qiagen, GmbH, Germany) real-time PCR machine following the manufacturer’s instructions. The melting curves were generated to ascertain of the amplification of a single product and the absence of primer dimmers. The cycle thresholds results were normalized to GAPDH as an endogenous gene. The expression levels were calculated as 2^ΔΔct^.

### Antibodies and western blotting

Polyclonal goat anti-human GRP78 (sc-1050), rabbit anti-human ATF4 (sc-200), rabbit anti-human CHOP (sc-200), rabbit anti-human GAPDH (sc-25778), rabbit anti-human MAP LC3 (sc-28266), horseradish peroxidase (HRP)-conjugated donkey anti-goat IgG (sc-2020) and HRP-conjugated goat anti-rabbit (sc-2004) antibodies were purchased from Santa Cruz Biotechnology (Dallas, USA). Western blotting was done as the procedure described previously [24,40].

### MTT proliferation and trypan blue survival assays

3-(4,5-dimethylthiazol-2-yl)-2,5-diphenyltetrazolium bromide assay and trypan blue cell staining was performed as described previously.[57]

### Statistical analysis

PCR primers were designed using IDT PrimerQuest software [58]. Image-based data were analyzed by ImageJ software [59]. The statistical significance was investigated by applying a two-tailed analysis of variance (ANOVA) using GraphPad Prism V6 software (GraphPad software Inc., La Jolla, USA). *P* ≤ 0.05 was considered as statistically significant. All the experiments were performed as minimum triplicate (N ≥ 3).

## Abbreviations

ANOVA: Analysis of variance
ATF6: Activating transcription factor 6
bZIP: Basic leucine zipper
C2Cer: C2-ceramide
CagA: Cytotoxin associated gene A
cagPAI: Cag pathogenicity island
DMSO: Dimethyl sulfoxide
ER: Endoplasmic reticulum
ERAD: Endoplasmic reticulum-associated degradation
Hp: Helicobacter pylori
HRP: Horseradish peroxidase
IDT: Integrated DNA Technology
IRE1α: Inositol-requiring kinase 1α
PBS: Phosphate buffered saline
PERK: PRKR-like ER kinase
T4SS: Type IV secretion system
Tm: Tunicamycin
UT: Untreated
XBP-1: X-box-binding protein.

## Acknowledgments

This study was granted by the Scientific and Technological Research Council of Turkey (TÜBİTAK; funding number 112S357) and Selçuk University Coordinatory of Scientific and Research Projects (BAP; funding number 10024925). The authors are thankful to TÜBİTAK and S.U. BAP for financial support of this study. The authors also thank Drs. Aydin Guzeloglu, Ercan Kurar for their helps and advices.

